# Assessing bimanual motor skills with optical neuroimaging

**DOI:** 10.1101/204305

**Authors:** Arun Nemani, Meryem Yucel, Uwe Kruger, Denise Gee, Clairice Cooper, Steven Schwaitzberg, Suvranu De, Xavier Intes

## Abstract

Measuring motor skill proficiency is critical for the certification of highly-skilled individuals in numerous fields. However, conventional measures use subjective metrics that often cannot distinguish between expertise levels. Here, we present an advanced optical neuroimaging methodology that can objectively and successfully classify subjects with different expertise levels associated with bimanual motor dexterity. The methodology was tested by assessing laparoscopic surgery skills within the framework of the fundamentals of laparoscopic surgery program, which is a pre-requisite for certification in general surgery. We demonstrate that optical-based metrics outperformed current metrics for surgical certification in classifying subjects with varying surgical expertise. Moreover, we report that optical neuroimaging allows for the successful classification of subjects during the acquisition of such skills.

---

Motor skills that involve bimanual motor coordination are essential in performing numerous tasks ranging from simple daily activities to complex motor actions performed by highly skilled individuals. Hence, metrics to assess motor task performance are critical in numerous fields including neuropathology and neurological recovery, surgical training and certification, and athletic performance^1–7^. In the vast majority of fields, however, current metrics are human-administered, subjective, and require significant personnel resources and time. Thus, there is critical need for more automated, analytical, and objective evaluation methods^4,8–11^. From a neuroscience perspective, bimanual task assessment provides insights into motor skill expertise, motor dysfunctions, interconnectivity between brain regions, and higher cognitive and executive functions, such as motor perception, motor action, and task multitasking^7,12^. Therefore, incorporating the underlying neurological responses in bimanual motor skill assessment is a logical step towards providing robust, objective metrics, which ultimately may lead to greatly improving our understanding of motor skill processes and facilitating bimanual-based task certification.

Among all non-invasive functional brain imaging techniques, functional near infrared spectroscopy (fNIRS) offers the unique ability to monitor and quantify fast functional brain activations over numerous cortical areas without constraining and interfering with bimanual task execution. Hence, fNIRS is a promising neuroimaging modality to study cortical brain activations but to date, only a very limited number of studies have been reported in regards to assessing fine surgical motor skills^13^. These exploratory studies have reported differentiation in functional cortical activations between groups with varying surgical motor skills ^13–17^. However, they suffer from recognized limitations ^13^ such as such as the lack of signal specificity between scalp and cortical hemodynamics ^18,19^, the lack of multivariate statistical approaches that leverage changes in functional brain activity across multiple brain regions, and benchmarking against established metrics. Hence, they have not impacted current practice of professional bimanual skill proficiency assessment. Here, we present a fNIRS-based optical neuroimaging methodology that overcomes all these shortcomings at once. For the first time in the field, we measure concurrently functional activations in the prefrontal cortex (PFC), the primary motor cortex (M1), and the supplementary motor area (SMA) to map the distributed brain functions associated with motor task strategy, motor task planning, and fine motor control in complex bimanual tasks^20–26^. Moreover, we increase the specificity of optical measurements to cortical tissue hemodynamics by regressing signals from scalp tissues^31,35,36^. Furthermore, we leverage changes in intraregional activation and interregional coupling of cerebral regions via multivariate statistical approaches to classify subjects according to motor skill levels. Finally, we compare our fNIRS based approach with currently employed metrics in surgical certification by assessing bimanual motor tasks that are a part of surgical training accreditation.

The performance of the reported optical neuroimaging methodology enables the objective assessment of complex bimanual motor skills as seen in laparoscopic surgery. Indeed, imaging distributed task-based functional responses demonstrated significant cortical activation differences between subjects with varying surgical expertise. By leveraging connected cerebral regions correlated to fine motor skills, we report increased specificity in discriminating surgical motor skills via fNIRS based metrics. For the first time, we show that our approach is significantly more accurate than currently established metrics employed for certification in general surgery, as reported via estimated misclassification errors. These results demonstrate that the combination of advanced fNIRS imaging with multivariate statistical approaches offers a practical and quantitative method to assess complex bimanual tasks. Topically, the reported optical neuroimaging methodology is well suited to provide quantitative and standardized metrics for bimanual skill-based professional certifications.

**Figure 1:**
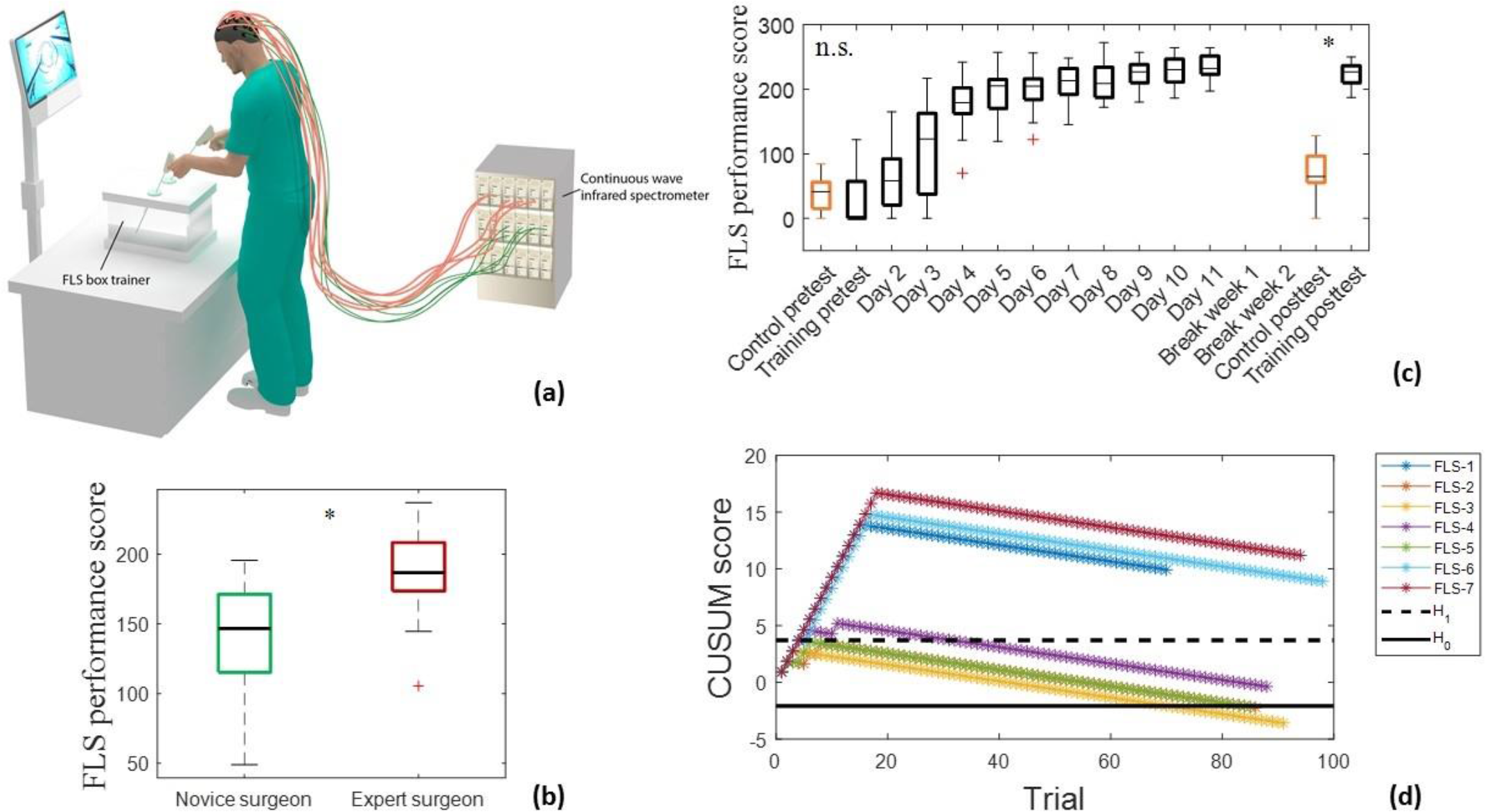
Bimanual motor task performance scores. **(a)** Schematic depicting the FLS box simulator where trainees perform the bimanual dexterity task. A continuous wave spectrometer is used to measure functional brain activation via raw fNIRS signals in real-time. **(b)** FLS performance scores for Novice surgeons (green) and Expert surgeons (maroon) where Expert surgeons significantly outperformed Novice surgeons. Two sample t-tests were used for statistical differentiation (^n.s.^ not significant, *p<0.05). **(c)** FLS performance scores for all training subjects (black) with respect to days trained compared to untrained Control subjects (orange). **(d)** CUSUM scores for each subject with respect to trials. The H_0_ threshold indicates that the probability of any given subject is mislabeled as a “Skilled trainee” is less than 0.05, and is subsequently labeled as a “Skilled trainee” subject. Results indicate that three subjects, FLS-2, FLS-3, and FLS-5 are labeled as “Skilled trainees”. The remaining subjects that do not cross the H_0_ line are labeled “Unskilled trainees”.

## RESULTS

### Surgical training task performance assessment

To demonstrate the potential of neuroimaging as an objective tool to assess bimanual task expertise, we selected a challenging bimanual pattern cutting task, which is part of the fundamentals of laparoscopic surgery (FLS) program. Demonstrating proficiency in the FLS is now required for certification in general surgery by the American Board of Surgery. For our study, we recruited a population with varying laparoscopic surgical expertise as defined via the FLS program and conventional professional nomenclature. The subjects were either classified into established skill levels, such as Novice surgeons (1^st^ – 3^rd^ year surgical residents), Expert surgeons (4^th^ – 5^th^ year residents and attending surgeons) or into trained medical students that are labeled as Skilled or Unskilled trainees (see **Supplemental Table 1**). The Control group constituted of medical students that underwent no training at all. Note that all groups were independent, i.e., each subject belonged to only one group. Each subject followed the official FLS pattern cutting task protocols. The experimental protocol followed by each cohort is provided in **Supplemental Figure 1**. The FLS performance scores were recorded for all subjects and the cumulative sum control chart (CUSUM) computed for the population following a training protocol. It is important to note that this study is the first to acquire FLS performance scores simultaneously with the neuroimaging data. The FLS scoring methodology was obtained with consent under a non-disclosure agreement from the FLS Committee. Thus, this study is the first one to report on direct comparisons of neuroimaging metrics and FLS scores for validation. In all cases, the FLS performance score were acquired simultaneously with the neuroimaging data.

Figure 1 (a) shows a schematic of the surgical trainer along with the fNIRS setup that is used to measure real-time cortical activation. A physical depiction of the setup is also provided in **Supplemental Figure 2**. Figure 1 (b) reports on the descriptive statistics of the FLS performance score for the Novice and Expert surgeons, where Experts significantly outperformed Novice surgeons (p<0.05). Similarly, the descriptive statistics of FLS performance scores over the whole training period are provided for all FLS task training subjects and untrained Control subjects in Figure 1 (c). Results indicate that there are no significant differences between the untrained Control subjects and training subjects on day 1 or the pretest (p>0.05). However, the trained FLS students significantly outperformed the untrained Control students on the final post-test, which follows a two-week break period post training (p<0.05). To provide insight at the subject level, Figure 1 (d) summarizes the CUSUM scores for each of the subjects with respect to trials performed. Trials that have a FLS performance score higher than 63 are considered a “success” and the respective CUSUM score is subtracted by 0.07 ^27,28^. Trials that have a performance score lower than 63 are considered a “failure” and the CUSUM score is added by 0.93 ^27,28^. Results indicate that three trained subjects (FLS 2, FLS 3, FLS 5) have passed the acceptable failure rate of 0.05 (H_0_) and thus are considered “Skilled” henceforth. The remaining trained four subjects (FLS 1, FLS 4, FLS 6, FLS 7) are considered “Unskilled” as they did not meet the FLS criteria for successful completion of the training program.

### Optical neuroimaging assessment of established surgical skill levels

To ascertain that our neuroimaging methodology can discriminate between established skill levels, we quantified the realtime hemodynamic activation over the PFC, M1, and SMA cortical regions while Novice and Expert surgeons performed the standardized FLS bimanual pattern cutting (PC) task ^27,29,30^, where typical hemodynamic responses are shown in **Supplemental Figure 5**. Figure 2 (a) depicts the spatial distribution of average changes in functional brain activation, as reported by Δ[HbO_2_] for all subjects in the surgical Novice and Expert groups. For the first time, significant differences were observed in all the PFC regions, the SMA, the left medial M1, and the right lateral M1 as depicted in Figure 2 (b). More precisely, Novice surgeons have significantly higher functional activation in the PFC regions (p<0.05) and significantly lower functional activation in the left medial M1 and SMA regions when compared to Expert surgeons. Habituation, the phenomenon where a response to a stimulus is gradually reduced due to repetition^31^, was not observed (p>0.05, data not shown).

While motor skill discrimination as reported via significant differences in the measurements from different cortical regions is typically central to neuroscience discovery studies, it does not provide insights into the utility of the data set to achieve robust classification based on quantitative metrics, such as accomplished during certification (*i.e.,* successfully pass a performance-based manual skills assessment). To quantify the performance accuracy of neuroimaging based classification of individuals in preset categories such as Novice surgeons (failed certification) and Expert surgeons (passed certification), we post-computed misclassification errors (MCEs) associated with current accredited FLS performance scores and with our neuroimaging method.

We employed a multivariate statistical method, namely linear discriminant analysis (LDA), to estimate the MCEs associated with the FLS and fNIRS based measurements. MCEs are defined as the probability that the first population is classified into the second population (MCE_12_) and the second population is classified into the first population (MCE_21_). Perfect classification is indicated by MCE = 0% and complete misclassification is indicated by MCE = 100%. Figure 2 (c) reports on these two misclassification errors for FLS performance scores and all combinations of fNIRS metrics for the classification of surgical Experts and Novices. Results indicate that subject classification is relatively poor when considering FLS performance scores only (MCE_12_=61% and MCE_21_=53%). On the other hand, neuroimaging based quantities provide lower errors (besides SMA only). Specifically, the combination of PFC, left medial M1 (LMM1) and the SMA leads to the overall lowest MCEs (MCE_12_=4.4% and MCE_21_=4.2%). Additionally, we provide the leave-one-out cross-validation results for the LDA classification models used for this data set, as seen in Figure 2 (d). This approach assesses the robustness of the LDA classification model, where each sample is systematically not used to build the LDA model and is treated independently. Results show that the combination of PFC+ LMM1 + SMA leads to the most robust and best performing data sets to build the classification model, as demonstrated by the fact that 100% of the samples in the leave-one-out cross-validation have MCEs<5%. The specific distributions of the classification results are shown in **Supplementary Figure 8a-b.** Furthermore, weights for each cortical region and their respective contribution to the total LDA model were also determined to show the correlation between different cortical regions on motor skill proficiency. The weights for left lateral PFC (0.58), medial PFC (0.23), right lateral PFC (0.29), left medial M1 (−0.70), SMA (0.14) contribute to the entire discriminant function with the norm of all the weights equal to 1.0. Three regions (left lateral PFC, right lateral PFC, and left medial M1) account for 96.18% of discriminant function indicating the preponderance of these regions for a robust and accurate subject classification.

**Figure 2:**
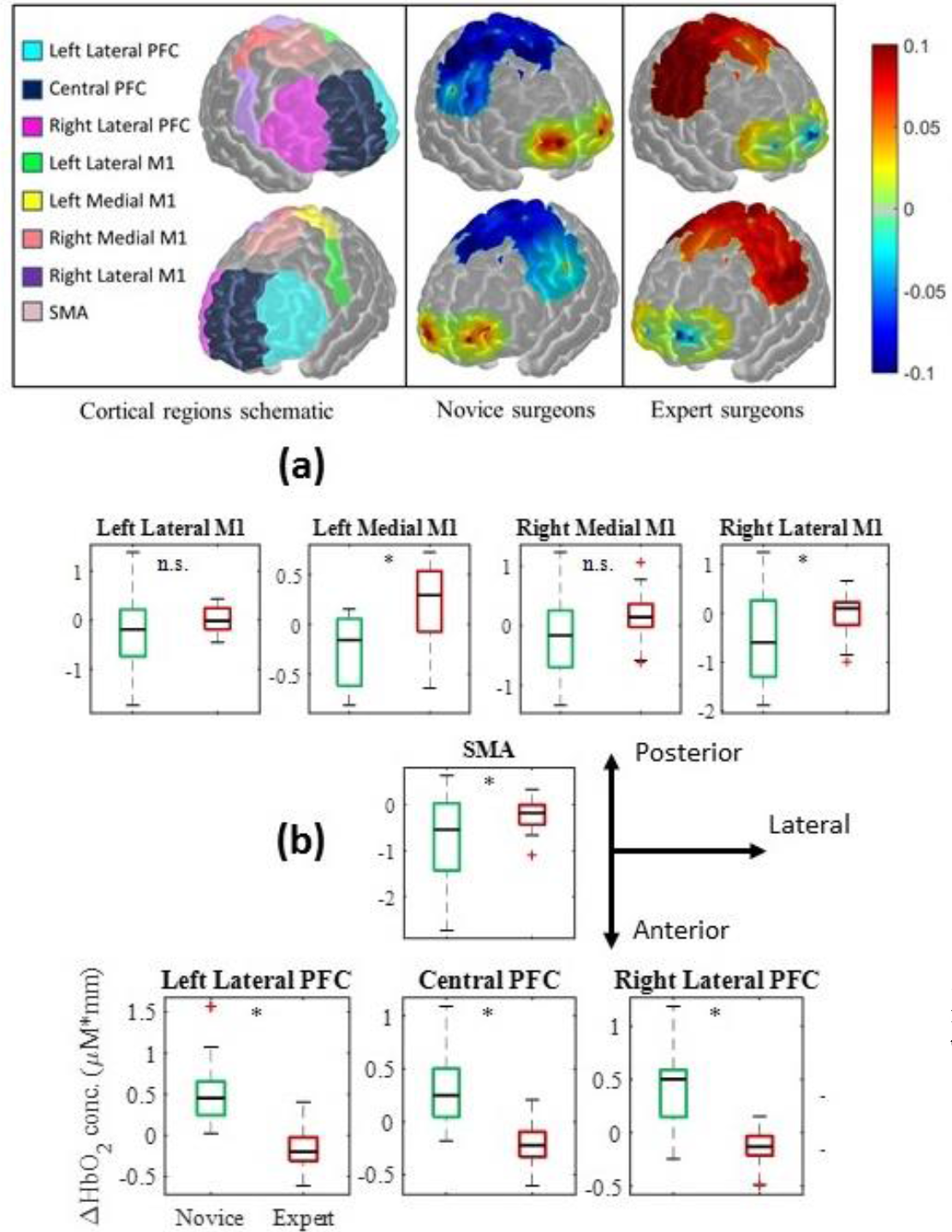
Differentiation and classification of motor skill between Novice and Expert surgeons. **(a)** Brain region labels are shown for prefrontal cortex (PFC), primary motor cortex (M1) and supplementary motor area (SMA) regions. Average functional activation for all subjects in the Novice and Expert surgeon groups are shown as spatial maps while subjects perform the FLS task. **(b)** Average changes in hemoglobin concentration during the FLS task duration with respect to specific brain regions for Novice (green) and Expert (maroon) surgeons. Two sample t-tests were used for statistical tests (^n.s.^ not significant, *p<0.05). **(c)** LDA classification results for FLS scores and all combinations of fNIRS metrics. **(d)** Leave-one-out cross-validation results show the ratio of samples that are below misclassification error rates of 0.05 for FLS scores and all other combinations of fNIRS metrics.

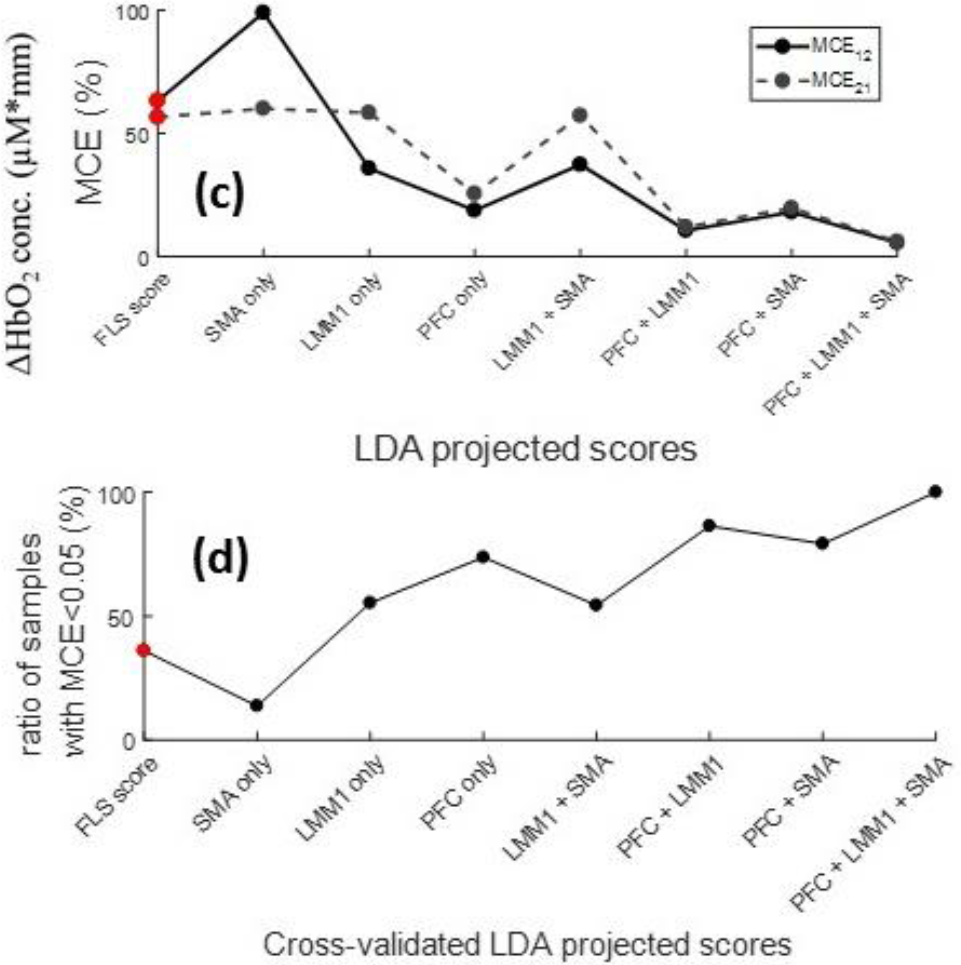

### Optical neuroimaging assessment of surgical skill level during training

Beyond determining skill levels of individuals compared to established groups, one key challenge in bimanual skill assessment and in laparoscopic surgery is the evaluation of bimanual motor skill acquisition during training. We applied our neuroimaging methodology to the FLS pattern cutting task over an eleven-day training period for inexperienced medical students. Based on the established FLS metrics currently employed in the field, the enrolled medical student population was divided into Skilled and Unskilled trainees at the completion of the training program as previously shown in Figure 1 (d). Additionally, five medical students with no prior experience in laparoscopic surgery were recruited as the Control group that underwent no training. Figure 3a shows a visual spatial map conveying the average cortical activation of all Skilled trainees or Unskilled trainees while performing the post-test (i.e. simulated certification exam). Like Expert vs Novice surgeons as shown in Figure 2a, Skilled trainees exhibit increased cortical activation in the left medial M1 and SMA and decreased PFC activation when compared to Unskilled trainees upon training completion and after a two-week break.

To provide a more global view of the training outcome, we present the descriptive statistics of functional activation between untrained Control students and all trained FLS students for pre-test (Day1) and post-test (final day after two-week break period) with respect to different brain regions in Figure 3 (b). Results indicate that there are no significant differences between the Control and all training students (Skilled and Unskilled trainees) at the onset of the training program (p>0.05). However, at the completion of the training and after a two weeks break period, both Skilled and Unskilled trainees exhibit a significantly lower functional activation in the left lateral and right lateral PFC compared to the untrained Control students (p<0.05). Furthermore, trained FLS students have significantly higher left medial M1 and SMA activation than untrained Control students during the post-test (p<0.05). These results reinforce the findings of the previous section regarding functional activation differences between Expert and Novice surgeons. To further stress the fact that our neuroimaging modality enables to provide a more granular view of training outcomes, we computed the MCEs for the three populations involved in this surgical training study (Control, Skilled and Unskilled trainees).

Figure 3c–d reports on the MCEs for each potential combination of medical student populations at different stages or end points of training. These MCEs were computed using the combined PFC, LMM1 and SMA brain functional optical measurements. The longitudinal MCEs of pretest populations versus odd days of training indicate that at the onset of the training, the populations could not be distinguished as reported by large inter-group misclassification errors, as shown in Figure 3 (c). However, after day 7, the Skilled trainee population demonstrated a significantly different neuroimaging distributed response compared to the first day of training, as demonstrated by very low intra-group misclassification errors.

**Figure 3:**
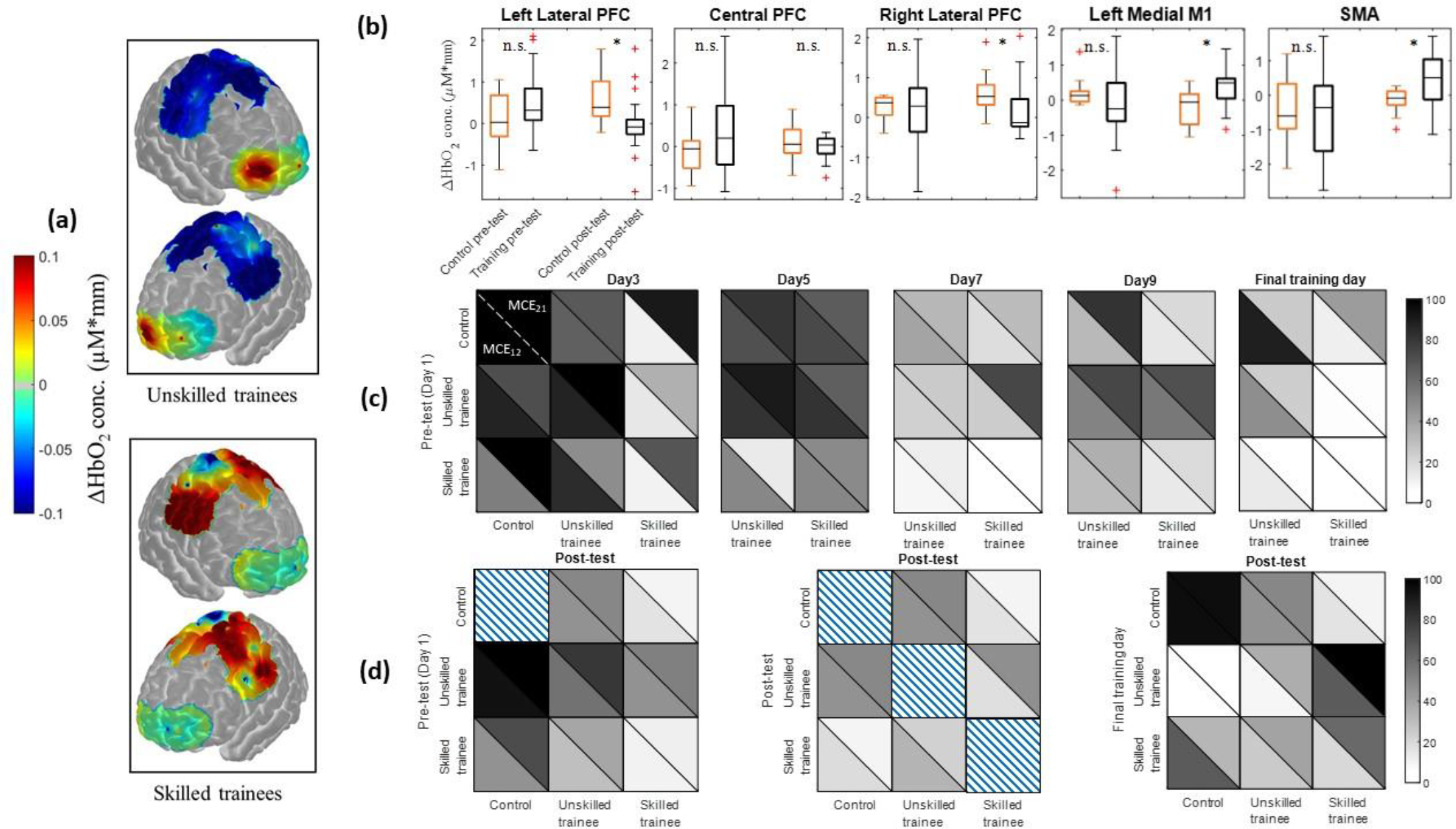
Differentiation and classification of motor skill between Control, Skilled, and Unskilled trainees. **(a)** Spatial maps of average functional activation for all subjects in each respective group during the FLS training task on the post-test day. **(b)** Average changes in hemoglobin concentration during stimulus duration with respect to specific brain regions for untrained Control subjects (orange) and all FLS training students (black). Two sample t-tests were used for statistical differentiation (^n.s.^ not significant, *p<0.05). Type I error is defined as 0.05 for all cases. **(c)** Inter and intra-group misclassification errors for each subject population (Control, Skilled and Unskilled trainees) with respect to training days. MCE_12_ and MCE_21_ values significantly decrease below 5% when classifying pre-test Skilled and Unskilled trainees on the final training day. Furthermore, misclassification errors are also low when classifying Skilled and Unskilled trainees on the final training day, along with Skilled trainees and untrained Control subjects. **(d)** Misclassification errors are reported for each combination of training groups (Control, Skilled, and Unskilled trainees) with respect to pre-test, post-test, and final training days. MCEs are substantially low when classifying Skilled trainees and Control subjects along with inter-Skilled trainee group classification. Unskilled trainees, however, showed high misclassification errors even when compared to Unskilled trainees and Control subjects during the post-test. As a measure of skill retention, classification models were also applied for all subject groups from the final training day to the post-test.

**Figure 4:**
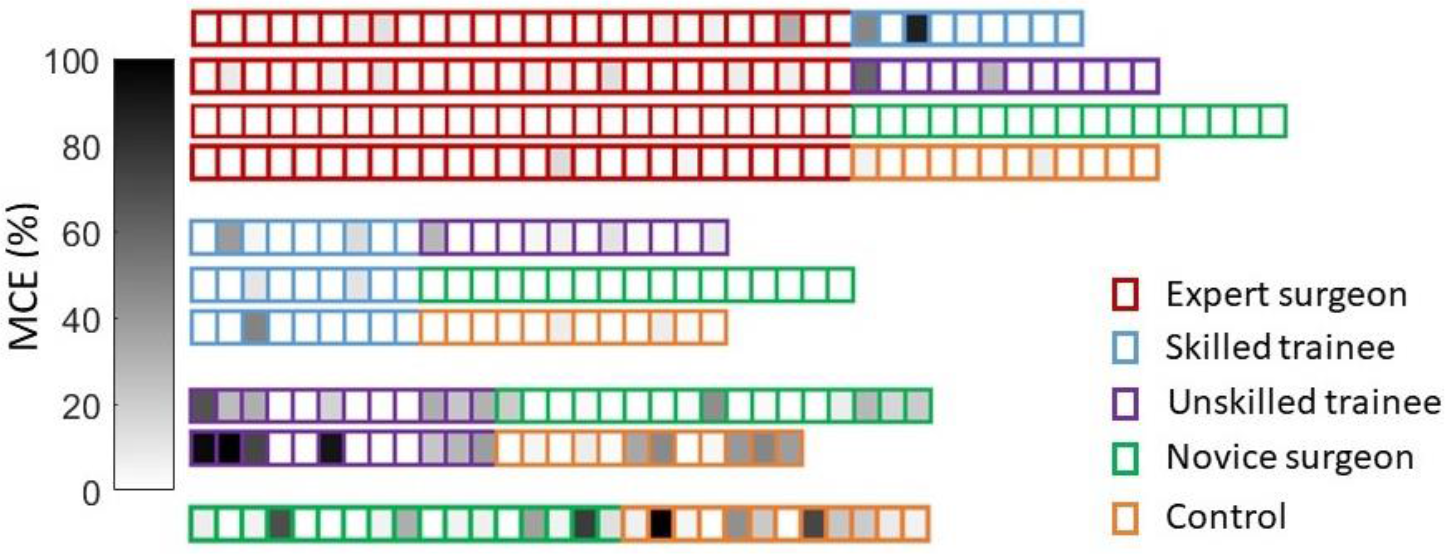
Cross-validation results for classification across all subjects with varying degree of motor skills. Each box represents one trial per expertise group during the post-test, where the shaded regions indicate the MCE if that given trial is removed from the classification model. Cross-validation results with their respective ratio of samples that are below misclassification error rates of 5% for Expert surgeons vs Skilled trainees (28/35 samples), Expert surgeons vs Unskilled trainees (29/ 38), Expert vs Novice surgeons (43/43), Expert surgeons vs untrained Control subjects (34/38), Skilled trainees vs Unskilled trainees (15/21), Skilled trainees vs Novice surgeons (24/26), Skilled trainees vs untrained Control subjects (18/21), Unskilled trainees vs Novice surgeons (16/29 samples), Unskilled trainees vs untrained Control subjects (11/24), and finally Novice surgeons vs untrained Control subjects (9/29).

Conversely, the Unskilled trainee population did not exhibit such marked trends. Even during the final training day (day 11), poor intra-group misclassification errors were observed for the Unskilled trainee population (MCE_12_ = 24% and MCE_21_ = 47%). In contrast, Skilled trainees on the final training day were completely classified from Skilled trainees on the pre-test, with MCE_12_ = 0% and MCE_21_ = 0%.

Similar results were observed when looking at the same intragroup misclassifications between the pre-test and post-test conditions, as shown in Figure 3 (d). Classification continues to remain poor for Unskilled trainees when comparing this population from the pre-test and the post-test, with MCE_12_ = 58% and MCE_21_ = 80%. Yet, Skilled trainees during the pre-test are successfully classified from Skilled trainees during the post-test, with MCE_12_ = 10% and MCE_21_ = 11%. While the Unskilled and Skilled trainee inter-groups were successfully classified at the end of the training session compared to the pre-test, the two populations did exhibit some intra-group overlap in their associated probability density function during the post-test. Of importance, both trainee populations did not exhibit marked differences between the final training day and post-test measurements as indicated by relatively high MCEs. Classification of Skilled trainees and Control subjects during the post-test also yielded in very low misclassification errors, whereas classification of Unskilled trainees and Control subjects still yielded in high misclassification errors, as shown in further detail in **Supplementary Figure 8c-d**. These cross-validated classification methods show that cortical activation has significantly changed for Skilled trainees during the post-test when compared to Skilled trainees on the pre-test or untrained Control subjects whereas Unskilled trainees do not exhibit such a marked trend.

### Classification of subjects with varying surgical expertise levels

For our neuroimaging based approach for motor skill differentiation to be formative, it is important to validate the classification models across all subject populations, especially since the studies associated with assessment of established skill levels and FLS training were performed independently in two different institutions. The subject population represents the full spectrum of laparoscopic surgical expertise, from Novices to certified attending surgeons, including Skilled and Unskilled medical student trainees. Regarding the number of procedures and associated level of expertise (at the completion of the training protocol), it is expected that the distribution in terms of surgical skills levels, from more proficient to less proficient is distributed as follow at the group level: Expert surgeons, Skilled trainees, Unskilled trainees, Novice surgeons, and Control.

Figure 4 shows the cross-validated classification model results comparing all subject population groups with varying expertise levels. Each box corresponds to a single trial for each expertise group, as shown via different colored borders. Shaded regions within each box indicate the MCE if that trial is removed from the classification model. For example, the first trial in cross-validation results for classifying Expert surgeons from Skilled trainees shows a MCE of 0%, as indicated by a white shade. However, the 29^th^ sample in the classification model, or the third trial in the Skilled trainee group, shows a MCE of 89% when removed from the classification model. The latter is an indication that the LDA classification model fails to reliably classify Experts and Skilled trainees if the third sample in the Skilled trainee group is removed.

First, we compare Expert surgeons with all other subject populations. Results indicate that Expert surgeons can be robustly classified with all subject populations, except for Skilled trainees where only 28/35 samples have MCEs less than 5%. Similarly, Skilled trainees can be successfully classified with Unskilled trainees, Novice surgeons, and Control subjects. Conversely, Unskilled trainees, Novice surgeons and Control subjects exhibit a poor inter-group classification as reported by multiple samples leading to high MCEs. Overall, these results indicate that the population with high expertise levels (Expert surgeons and Skilled trainees), can be robustly classified compared to groups that have not yet attain the required expertise levels as required by the FLS training program. “Non-certified” group including Novice surgeons, Unskilled trainees, and Control subjects, however, cannot be robustly classified between themselves.

## DISCUSSION

While there have been extensive efforts in the surgical community to confirm training effectiveness and validation of the FLS program ^29,30,32–34^, the surgical skill scoring component has received little attention and has garnered criticisms, such as subjectivity in scoring, inconsistencies in FLS score interpretations, and no correlation of patient injury reduction due to FLS certification ^30,33,35–39^. Despite the lack of rigorous evaluation of the FLS scoring methodology, the program has become the de-facto evaluation method for accreditation of skills required for general surgery ^37^. Given the high-stakes nature of surgical assessment in the FLS program and its implications on training future surgeons, there is a current gap in the rigorous validation of FLS scores as a robust and objective methodology ^37^. In this regard, previous studies have broached the concept of non-invasive brain imaging as a means for objectively assessing surgical skills ^13,14,17,40^. However, they suffer from methodological limitations that are now well-recognized by the fNIRS community, namely the contamination of superficial tissue, such as scalp, dura, or pia matter, in the recorded measurements^18,19^ (see **Supplemental Figure 4**). To highlight this point, results from the Expert and Novice surgeon cohort in this study were reprocessed without the regression of superficial tissue data and are provided in **Supplementary Figure 9a-c**. These results clearly demonstrate that previously reported fNIRS based metrics with the inclusion of superficial tissue responses can statistically differentiate surgical novices and experts ^13–16,40^, yet fail to classify subjects based on motor skill proficiency and perform as poorly as current surgical skill assessment metrics. In contrast, regressing shallow tissue hemodynamics from the optical measurements significantly reduces the false omission rate, where a surgical novice is mistakenly classified as an expert, to 0% whereas previous approaches still maintain false omission rates of 13-18% (see **Supplemental Table 2)**.

Beyond improving the robustness of optical measurements sensitivity to cortical activations, this work is also the first to measure functional activation in a multivariate fashion to determine critical cortical regions that are correlated to surgical motor skill differentiation and classification. More specifically, this is the first report of measuring functional activations in the PFC, M1, and SMA cortical regions that are putatively associated with motor task strategy, motor task planning, and fine motor control ^3,14,15,40–46^. Our results demonstrate that the inclusion of these cortical regions significantly improves the utility of fNIRS in assessing bimanual skills and can offer improved objective metrics over conventional FLS-based metrics currently used for certification in general surgery. Of importance, while using single regional readouts lead to enhanced population differentiation, the combination of the three above mentioned cortical regions provide excellent classification performances (for completeness, we also provide bivariate classification results using support vector machines in **Supplemental Figure 6-7**). Indeed, when combining measurements from these three brain regions, optical neuroimaging enables a remarkably robust classification of subjects based on their proven surgical skills levels, including novice, intermediate and expert skill levels. More precisely, our methodology allows for: (1) highly accurate classification of subjects with well-defined bimanual skills levels with better performance than currently employed metrics, (2) longitudinally assessing the acquisition of surgical skills during the FLS training program, and (3) performing robust classifications of populations recruited from multiple institutions with varying skill levels.

On a practical side, it is important to note that even if our methods leverage the most recent technical developments in the field of fNIRS, the instrumental and algorithmic platforms employed herein are readily available for wide-dissemination and use in surgical training facilities. Moreover, as more neuroscience-driven investigations focus on mapping distributed brain function, the positioning of the optodes (source or detector) on the subject scalp become increasingly challenging with extended spatial coverage. One key consideration is to ensure effective coupling is minimally affected by natural movements and not compromised by the subject’s hair. Hence, positioning of the optodes can be a lengthy process that is not suitable for professional environments that are time-constrained either by cost or throughput considerations. In this regard, our study identifies that the PFC, SMA and left medial M1 regions are sufficient for accurately assessing bimanual skill-based task execution. Thus, probe placement can be completed in a short amount of time without any impact on task execution, both critical factors for an acceptance of our surgical skill assessment methodology by the surgical community.

Beyond bimanual skills assessment and objective classification of individuals based on their skill levels, the work herein provides a sound foundation to further investigate the neurophysiology underlying bimanual skill acquisition and retention. Herein, we deliberately focused on reading the brain outputs as a mean to provide objective and quantitative measures of bimanual task execution without delving into the mechanistic understanding of the underlying physiology and functional connectivity. However, current neurophysiological knowledge supports the overall findings of our studies, namely increases in left medial M1 and SMA activation, and significant decreases in PFC activation across all groups with increasing motor task performance ^3,13–15,40–46^. It is also important to note that previous studies utilize motor tasks that are deliberately designed to decrease variability in studying cortical activation changes, such as finger tapping or simple visual or virtual based unimanual tasks. Conversely, the FLS task at hand is a complex bimanual task that involves visuospatial coordination, varying degrees of synchronicity between hands, motion frequency and range, and exerted forces on the surgical tools for task completion. Consequently, it is not feasible to ensure that each session replicates the same conditions and hence, the same cortical responses. Moreover, the cortical activations and interactions associated with the task planning and execution are dynamic by nature from expected explicit control in the early stages of learning to more implicit or automatic control in the later stages of motor learning. Thus, mapping the cortical networks and their dynamical changes associated with task execution and skill acquisition should be the next step.

Indeed, there is currently great interest in investigating dynamic functional connectivity (DFC) in neuroscience. Typically, DFC studies are conducted using fMRI, which is not appropriate for protocols requiring supine positions and/or non-elicited task execution. Recent studies have demonstrated that fNIRS is well positioned in such scenarios ^47–49^. We foresee that implementing such approaches in the context of bimanual skill assessment can lead to refined skill level assessment metrics as well as potentially provide predictive models of skill acquisition. For instance, composite cognitive metrics, possibly obtained by weighting regional cortical measurements using the LDA weights for best classification between Skilled and Unskilled trainees, could be central to developing tailored surgical training program for optimal skill acquisition and retention assessment (**Supplemental Figure 7)**. Furthermore, these methodologies can be easily applied to other fields including rehabilitation, brain computer interfaces (BCI), robotics, stroke and rehabilitation therapy ^50–52^. In summary, we believe this non-invasive imaging approach for objective quantification for complex bimanual motor skills will bring about a paradigm change in broad applications such as surgical certification and assessment, aviation training, and motor skill rehabilitation and therapy.

## Acknowledgement

This work was supported by funding provided by NIH/NIBIB grants 2R01EB005807, 5R01EB010037, 1R01EB009362, 1R01EB014305, R01EB019443. We would like to acknowledge David Boas and Jay Dubb from the Martinos Imaging Center for their significant support throughout this project. Finally, we would like to acknowledge Arthur “Buzz” DiMartino and his team at TechEn for their gracious support for the fNIRS hardware components.

## Author contributions

A.N. conceived the original idea. A.N., X.I., S.D and M.Y. designed the research study. D.G., C.C., S.S. and M.Y. contributed to study logistics, subject recruitment, and clinical expertise. A.N. performed the research studies. A.N., M.Y., U.K performed the data processing and analyses of results. A.N., M.Y., U.K., X.I., and S.D. interpreted the results. A.N. and X.I. wrote the manuscript and M.Y., U.K., and S.D. edited the manuscript. All authors discussed the conclusions and commented on the manuscript.

## Additional Information

Supplementary information is available in the online version of the paper. Reprints and permissions information is available online at www.nature.com/reprints Correspondence and requests for materials should be addressed to X.I.

## Data availability

The data, figures and other research findings in this study are available from the corresponding author on reasonable request

## Methods

The study was approved by the Institutional Review Board of Massachusetts General Hospital, University of Buffalo, and Rensselaer Polytechnic Institute.

## Hardware and equipment

We utilize a validated continuous-wave, 32-channel near-infrared spectrometer for this study, which delivered infrared light at 690nm and 830nm (CW6 system, TechEn Inc., MA, USA). The system employed eight long distance and eight short distance illumination fibers coupled to 16 detectors. The long-distance channels comprised all the measurements within a 30 – 40mm distance between the source and the detector, and the short distance channels comprised all the measurements within a ~8mm distance between the source and the detector. The short channels are limited to probing the superficial tissue layers, such as skin, bone, dura and pial surfaces, whereas the long channel probed both superficial layers and cortical surface. The probe design was assessed using Monte Carlo simulations and was characterized to have high sensitivity to functional changes in the PFC, M1, and SMA. A schematic of the geometric arrangement of probes are shown in **Supplementary Figure 2.**

## Participants and experimental design

17 surgeons and 13 medical students participated in this study. The minimum number of samples required for this study was determined *a priori* using power analysis according to the two-sample t-test comparing the means between two groups. Based on an initial pilot study, a conservative effect size (d=1.4) was chosen for the prefrontal and motor cortices. Furthermore, with 95% confidence interval, and a minimum power of 0.80, it was determined that a minimum of 8 samples were required per group, which was calculated by a statistical software G*Power^53^. The sample population was distributed within Novices (n=9, 1^st^ – 3^rd^ year residents with mean age 31 ± 2) and Experts (n=8, 4^th^ and 5^th^ year residents and attending surgeons with mean age 35 ± 6) surgeons. Subject demographics are listed in Table 1. To avoid any issues regarding hemisphere specific activation, only right-handed participants were selected. All participants were instructed on how to perform the task with standardized verbal instructions indicating the goal of the task and rules for task completion. The optical probes were positioned on the participant with great care to avoid any hair between the source/detector and scalp, as well as, robust coupling with the skin. The cap holding the fibers on the participant as well as the fibers did not hinder the participant’s movement during bimanual tasks. The participants were asked to perform the FLS pattern cutting task using a FLS certified simulator, where the goal is to use laparoscopic tools to cut a marked piece of gauze as quickly and as accurately as possible. The experiment for each participant consisted of a block design of rest and stimulus period (cutting task). The surgical cutting task was performed until completion or stopped after five minutes. Then a rest period of one minute was observed. The cycle of cutting task and rest periods was repeated five times per participant. The following measurements were recorded simultaneously for each participant during each trial: total task time, light intensity (raw NIRS data), and performance scores for the pattern cutting task based on the FLS metrics.

## NIRS post processing

Data processing was completed using the open source software HOMER2^54^, which is implemented in Matlab (Mathworks, Natick, MA). First, channels with signal quality outside of the range of 80dB to 140dB were excluded. The remaining raw optical signals (intensity at 690nm and 830nm) were converted into optical density using the modified Beer-Lambert law with a partial path-length factor of 6.4 (690nm) and 5.8 (830nm) ^55–57^. Motion artifacts and systemic physiology interference were corrected using recursive principal component analysis and low-pass filters ^54,58,59^. The filtered optical density data is used to derive the delta concentrations of oxy and deoxy-hemoglobin.

The short distance channels are regressed from the long-distance channels to remove any interference originating form superficial layers. This is achieved by using a consecutive sequence of Gaussian basis functions via ordinary least squares to regress scalp and dura activation data collected from the short separation fibers to create the hemodynamic response function (HRF) ^60,18,19^. Finally, the corresponding source and detector pairs for each source were averaged over each subject’s task completion time. The result is a scalar value for the change in oxy-hemoglobin according to different brain regions for all participants.

## Task performance metrics, statistical, and classification methods

The FLS scores were determined using the standardized FLS scoring metric formulation for the pattern cutting task based on time and error. This formulation is IP protected and was obtained with consent under a non-disclosure agreement from the FLS Committee, and hence its details cannot be reported in this paper. Descriptive and inferential statistics were performed using SPSS (IBM Inc., NY, USA). Two sample t-tests were used to determine statistically significant differences in functional activation between two groups. All box plots display median values (red bar) along with standard deviations. A confidence level of 95% was selected as the minimum required to reject the null hypothesis.

Linear discriminant analysis (LDA) was used to classify the populations based on their FLS scores and functional brain activation metrics. Prior to the analysis of LDA, all recorded metrics were first normalized, *i.e.* the sample mean and variance is 0 and 1. LDA determines the optimal vector, *v*, such that the projected metrics of two classes (ex. Novice and Expert surgeons) in the *v* direction has the highest separation between the classes with the lowest variance for each class ^61^. The resulting LDA scores are objectively compared for each class and the degree of separation objectively quantified as misclassification errors.

